# Audiovisual integration in the Mauthner cell enhances escape probability and reduces response latency

**DOI:** 10.1101/2021.08.19.456957

**Authors:** Nicolás Martorell, Violeta Medan

## Abstract

Fast and accurate threat detection is critically important for animal survival. Reducing perceptual ambiguity by integrating multiple sources of sensory information can enhance threat detection and reduce response latency. However, studies showing a direct link between behavioral correlates of multisensory integration and its underlying neural basis are rare. In fish, an explosive escape behavior known as C-start is driven by an identified neural circuit centered on the Mauthner cell. The Mauthner cell can trigger C-starts in response to visual and auditory stimuli allowing to investigate how multisensory integration in a single neuron affects behavioral outcome after threat detection. Here we demonstrate that in goldfish visual looms and brief auditory stimuli can be integrated to increase C-start probability and that this enhancement is inversely correlated to the saliency of the cues with weaker auditory cues producing a proportionally stronger multisensory effect. We also show that multisensory stimuli reduce response latency locked to the presentation of the auditory cue. Finally, we make a direct link between behavioral data and its underlying neural mechanism by reproducing empirical data with an integrate-and-fire computational model of the Mauthner cell.

## INTRODUCTION

When confronted with potential threats in their environment, animals have to decide and how to perform evasive movements to avoid harm. This is a powerful evolutionary force that led to the development of specialized sensory organs that extract qualitatively different information from a given event. The process of binding the different sensory signals associated to a single coherent event is called multisensory integration (1,2). Behaviorally, response improvement due to multisensory integration is often quantified by evaluating differences in the accuracy and speed of detection, localization and identification of stimuli (1,3). At the cellular level, integration performed by multisensory neurons is determined by synaptic convergence of sensory afferents belonging to different modalities, the neural operations that produce an “integrated” output and the interactions with other elements of the circuit or other brain areas (2). Multisensory integration becomes critically important for threat detection, when small reductions in sensory ambiguity can have a huge impact in the survival of the animal (4). After detecting a potential threat animals can perform a variety of protective behaviors. Depending on the perceived level of danger, animals might seek refuge, display freezing or alarm responses and when danger is extreme, fast escape behaviors (5–7). However, studies where behavioral correlates of multisensory integration can be directly tied to activity in an identified neuronal circuit are very rare and limited by the complexity and distribution of the neuronal networks involved.

Teleost fish can perform different evasive behaviors, the most explosive known as C-start fast escape. The C-start is initiated by a bilateral pair of reticulospinal neurons called Mauthner cells which determine the occurrence, latency and direction of the escape (8). Mauthner cells have two main dendrites: disynaptic auditory input arriving from the inner ear contacts de lateral dendrite of the Mauthner cell (9–11) while polysynaptic visual input coming from the optic tectum contacts the ventral dendrite of the cell (12,13). Both auditory and visual stimuli can activate the Mauthner cell with a probability and response latency that are function of the stimulus salience (11,14,15). Auditory salience depends on the amplitude of the sound wave while visual salience has been tied to stimulus dynamics and contrast (14,16–19). The temporal structure of auditory or visual stimuli that consistently trigger C-starts differs (17,20). Auditory C-starts are triggered by intense and brief auditory pips while visual C-start responses are typically evoked by fast expanding dark disks (14,16,21). The one-to-one relationship between a C-start and firing in the Mauthner cell offers a unique opportunity to study how multisensory integration in a single neuron impact on a fast escape behavior.

Here we demonstrate that in goldfish auditory and visual stimuli can be integrated to enhance C-start probability and reduce response latency. We also show that an integrate-and-fire computational model of the Mauthner cell reproduces fish responses to multisensory stimuli, making a direct link between behavioral data and its underlying neural mechanism.

## METHODS

### Animals

One hundred and ninety adult goldfish (*Carassius auratus*) of both sexes, 7–10 cm of standard body length were used in this study. Fish were purchased from local aquaria (Daniel Corralelo, Buenos Aires, Argentina) and allowed to acclimate for at least a week after transport. Fish were kept in rectangular glass holding tanks (30 x 60 x 30 cm; 95 l) in groups of 10. Tanks were supplied with filtered and dechlorinated water and maintained at 18°C. Ambient light was set to a 12-h light/dark photoperiod. Animals were fed floating pellets (Sera, Germany) five times a week.

All procedures were performed in accordance with the guidelines and regulations of the Institutional Animal Care and Use Committee of Facultad de Ciencias Exactas y Naturales, Universidad de Buenos Aires (protocol #70).

### Experimental Setup

Goldfish were tested in a rectangular experimental tank (80 cm length, 70 cm width, and 15 cm height) standing on an anti-vibration table with its external walls covered with black opaque cardboard to avoid external visual stimulation. A cylindrical enclosure (60 cm diameter, 13 cm height) made of clear acetate was placed inside the tank to confine fish to the experimental arena (Figure 1A). The tank was filled with filtered dechlorinated water up to a height of 12 cm. Auditory stimuli were produced by an underwater loudspeaker (UW-30, University Sound, Buchanan, MI) placed outside the experimental arena and supported within a 6 cm thick layer of foam. Visual stimuli were produced by a video projector (Epson Powerlite S6+, 60 Hz) secured 130 cm above the tank. The tank was covered by a lid made of white wax paper sheet, which served as a projection screen for the visual stimuli. The bottom of the tank was transparent, allowing for simultaneous video recording of fish behavior and stimulus presentation at 240 fps (432×320 pixels, Casio EX ZR100, Tokyo, Japan). A green LED located below the experimental tank (hidden from animal’s view) was used to allow the camera to record the time of auditory stimulus onset. Experiments were made in a silent room with ceiling lights off. Presentation of visual and auditory stimuli as well as camera acquisition and LED activation was controlled by MATLAB (The Mathworks). Triggering of video acquisition occurred simultaneously with visual stimulus onset and stopped 8.25 s after the end of visual stimulation.

**Figure 1.**
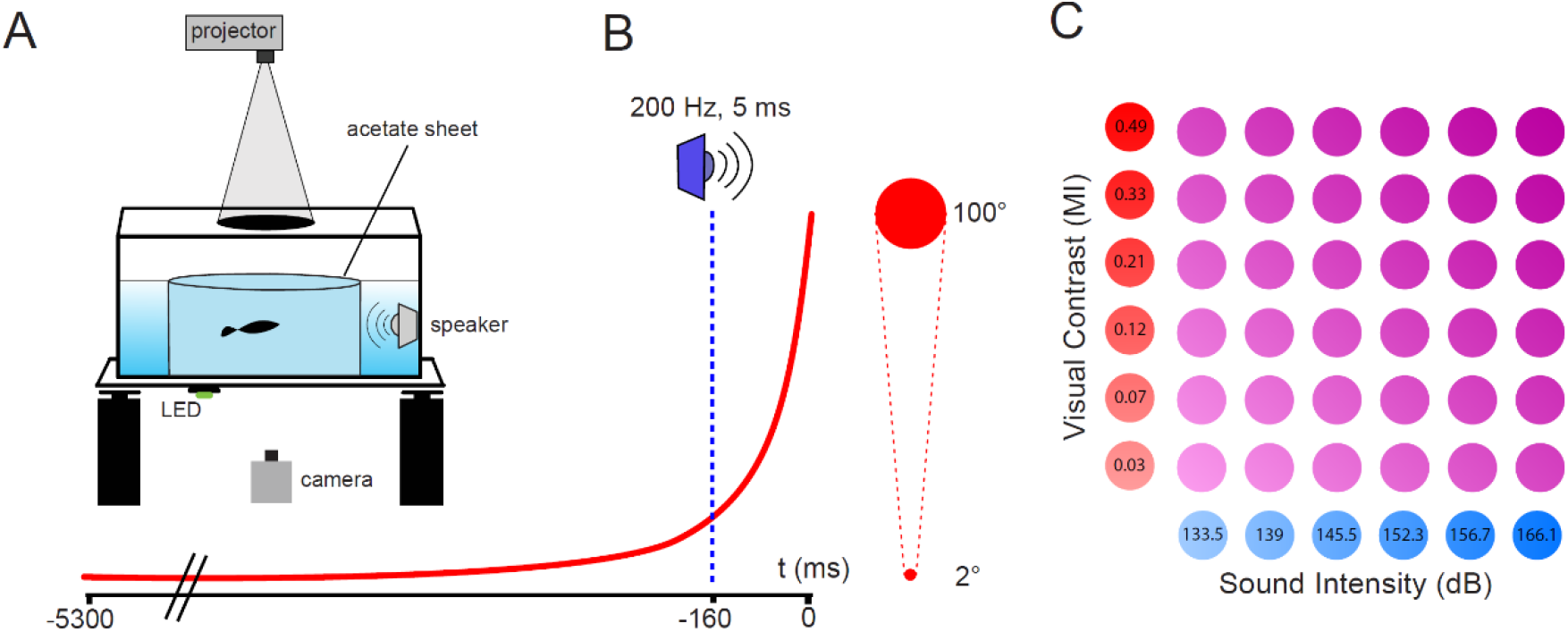
Experimental Setup and Stimulus Design. (A) Behavioral setup. Computer-generated visual and auditory stimuli were delivered through a video projector and a water-proof speaker, respectively. A clear acetate sheet delimited the experimental arena. An LED was turned on synchronously with auditory stimulus onset while a high speed camera recorded fish behavior, visual stimuli and the state of the LED for the duration of each trial. (B) The visual loom expanded from a subtended angle of 2° to 100° in 5.3 s (L/V=0.192). When visual and auditory stimuli were combined, the auditory stimulus onset preceded the end of visual expansion by 160 ms. (C) Six intensities of visual stimuli (Michelson Index, MI) and 6 intensities of auditory stimuli (amplitude in dB, re 1μPa) were combined into 36 different multisensory stimuli, thus creating a 2D space of multisensory intensities.

### Stimuli

Computer-generated dark disks that expand over a lighter background (loom) efficiently elicit C-start escapes (14,16,21). In our study, looming stimuli expanded exponentially from a diameter of 0.27 cm until reaching a diameter of 18.15 cm 5.3 s later. The subtended angle, as measured from the center of the tank, grew from 2.06° to 100.84° (Figure 1B). Looming stimulus can also be defined by the size of the approaching object of equal width and height (L), the approach velocity (V), and the apparent distance covered. The combined term L/V is an indicator of the expansion of the subtended angle with time and is used to allow comparison to looming-evoked responses in other animals. The looming used here yield an L/V=0.192. We chose a fast expanding loom as it has been shown that low L/V values preferentially recruit fast escapes triggered by the Mauthner cell whereas looms of higher L/V values might recruit networks driving responses of longer latency (17,20,22).

To obtain visual stimuli of different salience, we modified the grayscale value of the disk stimulus (RGB values between 70 and 118) while keeping the background constant (RGB value of 120), obtaining different contrasts (19,23). To characterize the luminance of each component of our stimuli, we used the irradiance sensor (J1812) of a Tektronix J17 photometer (Wilsonville, Oregon, MI, USA) positioned in the center of the tank while projecting images on the wax paper lid. During these measurements, all pixels in the screen were set to the RGB value that we were currently testing. Using these irradiance values, we determined the contrast for each stimulus, calculated as the Michelson index (MI), where contrast is defined as (I_disk_ - I_background_) / (I_disk_ + I_background_), and I_disk_ and I_background_ refer to the irradiance of the expanding disk and the background, respectively. We used six different visual stimuli, characterized by Michelson indices of 0.03, 0.07, 0.12, 0.21, 0.33 and 0.49. Auditory stimuli consisted of a single cycle of a 200 Hz sine wave (5 ms in duration), whose amplitude was changed to modify its salience. We used six different auditory stimuli of amplitudes of 133.5, 139, 145.5, 152.3, 156.7 and 166.1 dB re. 1 μPa when recorded with a hydrophone (Sensor SQ34) placed 10 cm away from the speaker. Intensity of auditory and visual stimuli was selected to obtain a range of unimodal C-start response probabilities spanning from about 0 to 80% (see below).

We analyzed 12 unisensory stimuli (6 visual, 6 auditory) as wells as all 36 multisensory combinations of stimuli (Figure 1C). When visual and auditory stimuli were combined (multisensory presentations), the auditory component onset occurred 160 ms before the end of visual expansion, unless otherwise specified (Figure 1B).

### Stimulation protocol

Individual fish were placed in the experimental tank and allowed to acclimate for 15 min. The animals then received between 9 and 13 stimuli that included at least one unimodal visual (V), one unimodal auditory (A) and 7 to 9 multisensory (V + A) stimuli. From the 36 possible multimodal stimuli each animal was tested with a subset that included low, medium and high intensity combinations. Stimulus presentation was randomized and intertrial interval set to 4 min to minimize the effect of habituation. However, a slight decrease in response probability was observed, both for C-start responses (21% of decrease by trial 13, Binomial GLM, p = 0.004) and alarm responses (4.6% of decrease by trial 13, Binomial GLM, p = 0.002, Supp. Figure 1A).

We also found a strong thigmotactic behavior, with 95% of the fish remaining within 15 cm from the enclosure walls (distances to the center of the arena were not consistent with a uniform distribution: Chi-Square Test of Homogeneity, p < 0.0001; mean distance to center ±standard deviation were of 29.4 cm ± 4.8 cm, Supp. Figure 1B, C). As visual looms were projected in the center of the arena, fish position affected the stimulus’ subtended angle perceived at the retina which could in turn affect C-start response probability. Indeed, C-start probability was higher for animals closer to the center of the arena (and to the expanding loom) and decreased as the animal’s position approached the enclosure walls (Supp. Figure 1D). However, this dependence could have been related to the position respect to the center of the arena itself and not to the fish’s relative position to the center of the loom expansion. To test that this was not the case, the visual expansion was centered on one extreme of the enclosure, and thus did not coincide with the center of the tank. When presented this way, C-start probability was uncorrelated with distance to the center of the arena (binomial GLM, p = 0.911). Throughout all trials, C-start probability was (negatively) correlated with the distance between the fish and the center of the looming expansion (binomial GLM, p < 0.0001, Supp. Figure 1D). As for auditory stimuli, distance to the speaker was not correlated with response probability (binomial GLM, p = 0.719, Supp. Figure 1E).

### C-start Escape and Alarm Responses

Videos were analyzed offline using VirtualDub (version 1.10.4, www.virtualdub.org) and custom-made code built upon the OpenCV library in Python (opencv.org). For trials which included a visual stimulus, the last frame of expansion of the looming was set as time point 0 ms. Therefore, C-start responses occurring before the end of visual expansion have a negative response time, whereas those occurring after rendered a positive response time (Figure 1B). For auditory only trials, response time was considered from the start of the auditory stimulus, which in the videos was recorded by the activation of an LED not visible to the fish. Animals which never performed a C-start response were excluded from analysis (N=10/190). Videos were also inspected and scored by two independent observers to analyze the occurrence of C-start escapes or other behaviors suggesting increased arousal or alarm. Alarm responses consist on a variety of subtle but robust motor reactions including accelerating or decelerating swimming, darting (a single fast acceleration in one direction with the use of the caudal fin), erratic movements/zigzagging (representing fast acceleration bouts in rapid succession), rapid abduction of fins with no body displacement and freezing (which consists of a complete cessation of movement, except for gills and eyes) (5,6,24). An alarm response was computed when scoring of occurrence and description of behavior matched for both observers.

### Data Analysis

In order to analyze the effect of multisensory integration for different treatments, we first asked what would be the expected response probability for a combination of visual and auditory stimuli if there was no integration (i.e. if visually-evoked responses and auditory-evoked responses were completely independent). Given a response probability for a visual stimulus P(V), and a response probability for an auditory stimulus P(A), the Expected Response Probability (ERP) in the absence of integration can be calculated as P(V or A) = P(V) + P(A) - P(V) ^*^ P(A). We compared the ERP with an Observed Response Probability (ORP) for each multisensory combination of stimuli V and A. The relative difference between ORP and ERP is a measure of the effect of integration over response probability. We define an Integration Coefficient (IC) as 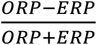. This value ranges from -1 to 1, it is 0 if integration has no effect over response probability, it is positive if integration increases response probability and it is negative if integration decreases response probability. We calculated the IC for each of the 36 multisensory combinations of stimuli.

### Statistical Analysis

R (version 4.0.2, www.r-project.org) and RStudio (version 1.1.456) were used for statistical analysis. A significance level of α = 0.05 was used throughout the study. Effects of explanatory variables over response variables were assessed using Generalized Linear Models (binomial GLMs in the case of binary response variables, gaussian GLMs in the case of continuous response variables). Sample size is denoted by *N* when it refers to the number of animals or *n* when it refers to the number of trials. When working with probability data, error bars represent standard error of a proportion.

### Computational Model

We used NEST 2.20.0 (25) running on Python 3.8.3 to create a computational model of the Mauthner Cell (Figure 8A) using a leaky Integrate-and-Fire neuron model with exponential post synaptic currents (iaf_psc_exp). This model follows a differential equation that can be expressed as 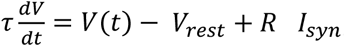, where *τ* is the time constant, V(t) is the membrane potential, V_rest_ is the resting potential, R_in_ is the input resistance and I_syn_ is the sum of the input currents. The parameters of this model were adjusted to match reported intrinsic properties of the Mauthner cell. Resting membrane potential was set to -80.0 mV and so was the reset value for the potential after a spike. Firing threshold was set to -65.0 mV, C to 2500 pF/cm2 and the time constant to 0.5 ms (13,26).

The Mauthner Cell receives excitatory mechanosensory inputs through its lateral dendrite and excitatory visual inputs through its ventral dendrite which are integrated at the soma. We modelled these two types of inputs while retaining stimulus characteristics from our experiments. Auditory stimuli were designed as 20 ms square pulses of current, as an approximation of recordings of Mauthner Cells during similar auditory stimulation (11,27). We designed 6 auditory stimuli with intensities that ranged between 75 nA and 250 nA. The intensities were adjusted to reproduce the observed unisensory auditory probability. Similarly, 6 visual stimuli were designed as increasing currents which followed the equation 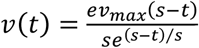, where *e* is Euler’s constant, *t* is time, *v*_*max*_ the maximum value of the “visual” input current and *s* is a parameter which controls the slope of the curve. *v*_*max*_ was varied to modify stimulus intensity, and ranged between 90 nA and 220 nA to match experimental C-start probabilities of unisensory visual trials. The value of *s* was picked randomly for each simulated trial from a Gamma distribution with a mean of 200 and a standard deviation of 150. This generates a family of curves which both resemble the temporal dynamic of our looming stimuli and of previous recordings of goldfish Mauthner Cells during looming stimulation (14). To add variability to the model, *v*_*max*_ and *a*_*max*_ (maximum value of “auditory” input) were multiplied by a number (R_1_ and R_2_, respectively) picked from a uniform random distribution in the interval (0; 1] before the beginning of each trial. Thus, reported stimulus intensities for auditory or visual stimuli represent the maximum possible input current (R_i_=1). The neuron received auditory and visual inputs through two distinct excitatory synapses with weights equal to 1.

For each combination of *v*_*max*_ and *a*_*max*_ the simulation was run 200 times. Although experimental trials lasted 5.3 s, behavioral responses were only observed in the last 1000 ms of visual stimulation, therefore computational rounds were only simulated for 1300 ms. The end of visual exponential increase was set to 300 ms before the end of the trial, and the auditory square pulse was presented 160 ms before the end of the visual input (unless otherwise specified).

We calculated response probability for a given stimulus as the proportion of trials where threshold was reached. Response time was recorded as the time elapsed between the threshold was crossed and the end of visual expansion (or to auditory stimulus onset in unisensory auditory trials).

## RESULTS

### Risk assessment is affected by prior motor state

Fish reacted to visual looms and auditory pips (Figure 1) with a variety of motor behaviors that were not limited to fast C-starts but included a range of slower escapes and alarm responses such as zigzagging, backward swimming, darting and freezing (5,6,20,28). As risk assessment and the resulting behavioral decision can be influenced by the animal’s state *prior* to sensory stimulation, we analyzed the fish motor behavior immediately before and after stimulation (7).

Combined responses of 180 animals to auditory, visual or multisensory stimuli showed that behavior expressed after stimulation was dependent on the previous motor state (χ2 Test of homogeneity, p < 0.0001). In about two thirds of all trials (63.3%, n=1003) fish were either actively swimming or remained still but beating fins while in the remaining trials fish were freezing before the stimulation begun (36.7%, n=580, Figure 2). Freezing had a strong impact on the behavioral outcome, in only 24% of trials in which animals were freezing a C-start was observed while the rest remained freezing (1% of trials individuals performed an alarm responses). In contrast, in trials from non-freezing animals 54% percent performed a C-start while 17% responded with an alarm behavior. Conversely, almost 80% of all C-starts were performed by animals that were not freezing at the moment of stimulation. Therefore, although freezing does not abolished completely the possibility of performing a C-start, it sharply reduces its probability.

**Figure 2.**
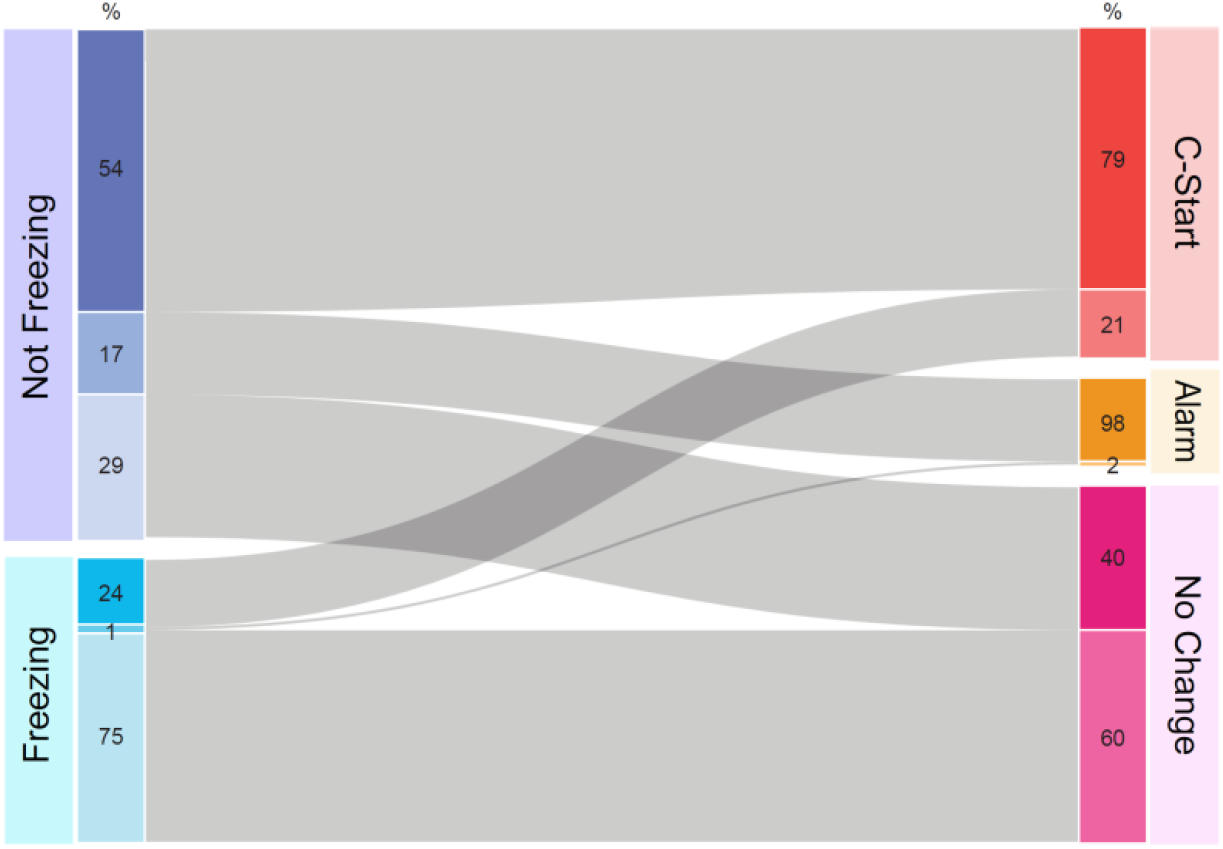
Risk assessment is affected by prior motor state. Alluvial diagram of fish motor behavior in each trial before and after stimulation. The diagram shows the motor state before (Not Freezing, n=1003 or Freezing, n=580) and after sensory stimulation (C-start, n=675; Alarm, n=176; No change in motor behavior, n=732) combining all stimulus conditions. Each motor state is further divided by its “future” or “past” action (numbers within colored boxes). For example, 54% of non-freezing animals executed a C-start, while 60% of animals that did not changed their motor behavior were freezing before stimulation.

To further explore the influence of previous motor activity on C-start occurrence we calculated swimming velocity during a 420 ms window, prior to auditory stimulation or 1.5 s before the end of visual expansion, since by this time no fish had responded (Supp. Figure 2 inset, grey area). Most animals swam at low velocities (<20 mm/s) and swimming velocity distribution did not differ between animals that subsequently responded with a C-start or not (Supp. Figure 2A, binomial GLM, p = 0.686). This suggests that, although freezing reduces C-start probability, swimming does not enhance *per se* response probability. In fact, still animals (fish with swimming velocity lower than 0.1 cm/s) had a C-start response probability three times larger than those freezing (Supp. Figure 2B, 56% for non-freezing animals (n = 267), 18% for freezing animals (n = 401), binomial GLM, p < 0.0001).

### Unisensory stimulus intensity modulates C-start response in non-freezing animals

To establish how varying unisensory stimulus salience affects risk assessment we evaluated the fish response to increasing loom contrast or sound intensity (Figure 3). In non-freezing animals, C-start response probability increased with stimulus intensity, covering a similar range in visual and auditory presentations (Figure 3A-B, visual: from 10% to 71%, binomial GLM, p < 0.0001; auditory: 0% to 76%, binomial GLM, p < 0.0001). Unisensory stimuli produced alarm responses (Figure 3C-D) although C-start probability was consistently higher than alarm probability, both for visual stimuli (red, Figure 3A vs. 3C, Fisher’s Exact Test, p < 0.0001) and auditory stimuli (blue, Figure 3B vs. 3D, Fisher’s Exact Test, p = 0.0008). Consistent with previous work from our lab (19), alarm response probability was not dependent on stimulus intensity, neither for visual stimuli (Figure 3C, binomial GLM for non-freezing trials, p = 0.868) nor for auditory stimuli (Figure 3D, binomial GLM for non-freezing trials, p = 0.141; binomial GLM for freezing trials, p = 0.164).

**Figure 3.**
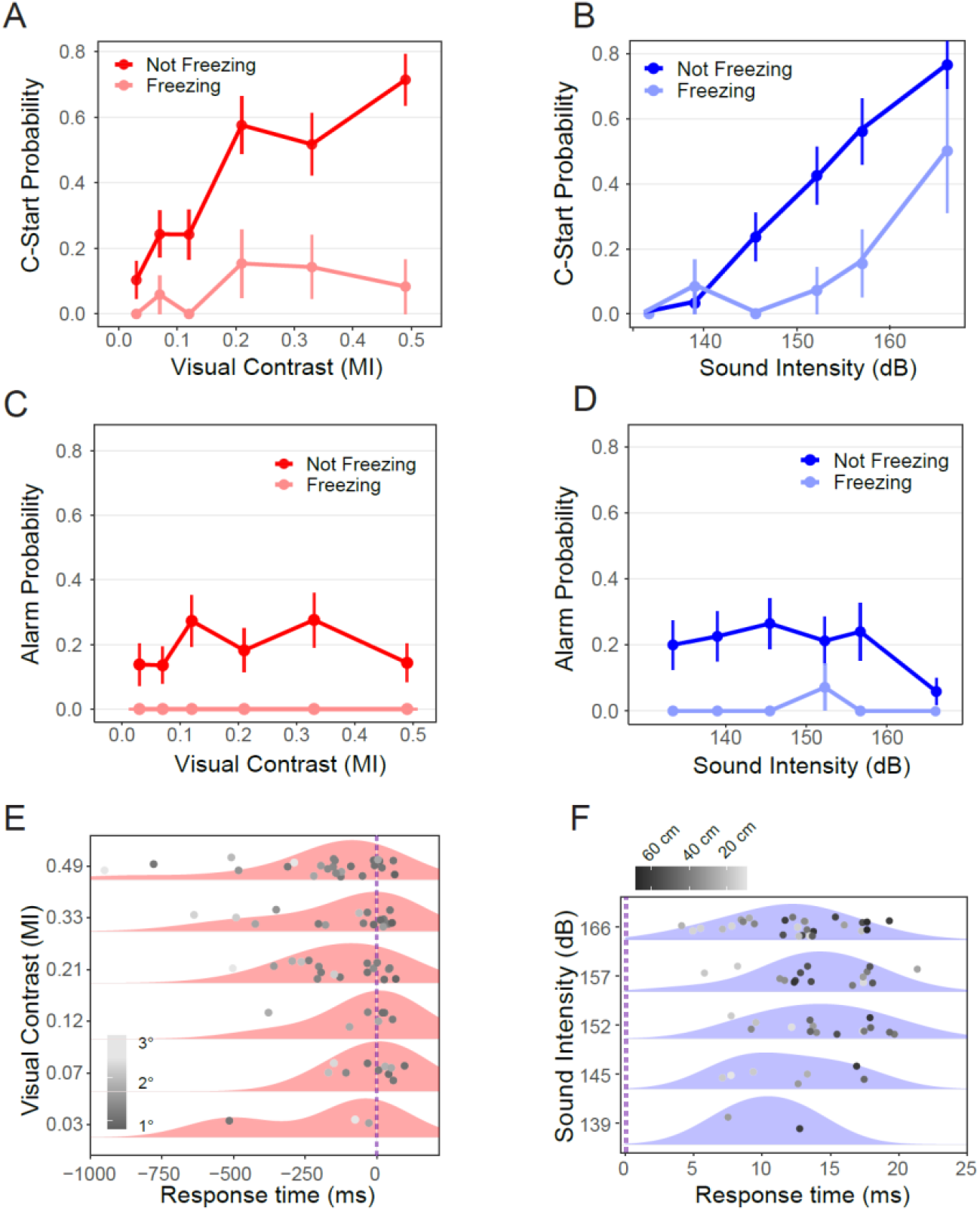
Unisensory stimulus intensity modulates C-start response in non-freezing animals. Unisensory C-start probability as a function of visual contrast (A) or auditory intensity (B) for non-freezing and freezing fish. Number of trials vary between n=29-37 for non-freezing and n= 12-19 for freezing trials. Alarm probability as a function of intensity for visual (C) or auditory (D) unisensory stimuli for non-freezing and freezing fish. Number of trials vary between n= 25-34 for non-freezing and between n=8-14 for freezing trials. In A-D bars represent standard error for the proportion. (E) Individual response times and density distributions for C-starts grouped by visual contrast (n = 83). Dots are shaded according to the looming’s subtended angle (°) from the perspective of the animal, measured 1250 ms before the end of expansion. Temporal scale is zeroed at the end of the loom expansion (dashed line). (F) Individual response times and density distributions for C-starts grouped by sound intensity (n = 70). The lowest intensity yielded no responses. Dots are shaded according to the animal’s distance to the speaker (cm), 1 s before stimulus onset. Temporal scale is zeroed at the onset of auditory stimulation (dashed line). Shaded red or blue areas show density distributions of response times computed using a Gaussian kernel. Vertical scatter was added for clarity in E and F.

Freezing animals had a lower C-start response probability across all visual or sound intensities (Figure 3A red vs. light red, binomial GLM, p < 0.0001; Figure 3B, blue vs. light blue, binomial GLM, p < 0.0001) did not performed alarm responses during visual stimulation (Figure 3C) and only once to sound stimuli (Figure 3D, n = 1/71). Finally, while C-start probability is independent of visual contrast in freezing animals (Fig. 3A, binomial GLM, p = 0.189), there is a significant correlation with auditory intensity (Fig. 3B, binomial GLM, p = 0.003), although this correlation is only observed at larger amplitudes.

### Stimulus intensity modulates response time

We next analyzed if C-start response time was affected by stimulus intensity. Our rationale was that as stimulus salience increases, the decision threshold should be reached faster. Responses to visual looms spanned within a wide time window, from 960 ms before to 100 ms after the end of the visual stimulation (Figure 3E). However, as contrast grows there is a left-shift in the response time density distribution, i.e. the proportion of early responses increases. Statistical analysis confirmed that higher contrast looms evoked C-starts with a shorter response time (gaussian GLM, p = 0.026). Response time was also influenced by freezing and the position of the animal in the experimental arena at the time of stimulation (supp. Figure 1B). Freezing delayed C-start onset (gaussian GLM, p = 0.005), and animals which experienced larger initial subtended angles (lighter grey points, Figure 3E) exhibited shorter response times (gaussian GLM, p = 0.002).

In contrast to the wide temporal distribution of visual responses, auditory C-starts occurred between 4 and 21 ms after sound presentation (Figure 3F). The density distributions of auditory responses peaked between 10-15 ms, a value well within that reported in the literature (29) and were not shifted by sound intensity (gaussian GLM, p = 0.737) nor were affected by freezing (gaussian GLM, p = 0.122). However, fish closer to the sound source responded earlier (gaussian GLM, p < 0.0001, lighter grey points, Figure 3F) (27). Animals located between 0-30 cm from the speaker had a mean response time of 10.64 +/- 0.82 ms while those between 31 and 60 cm had a response time of 14.75 +/- 0.67 ms (Two Sample T-Test, p = 0.0003).

### Multisensory integration enhances C-start probability

We also wanted to analyze uni vs. multisensory processing to assess how animals integrate multisensory information during threat detection. To this aim we designed sensory stimuli that mimic some aspects of the approach of a predator from above (e.g. a bird), in direct collision course (looming stimulus). As the aerial predator finally breaks the surface of the water, a loud splash (auditory pulse) briefly precedes the end of the looming expansion (Figure 1B). Besides being an ecologically relevant situation, this stimulus structure allowed us to analyze specifically as the effect of sudden addition of information (sound) after a period of gradual evidence accumulation (loom) in a potentially dangerous scenario. Combination of 6 levels of visual contrast and 6 levels of sound intensity resulted in 36 distinct multisensory stimuli, each with its unique combination of visual contrast and sound amplitude (Figure 1C). Figures 4 shows the C-start probability for each combination for non-freezing (A) and freezing (B) animals. In line with the unisensory effects reveals that C-start probability increases with contrast, sound amplitude while freezing reduces response probability (binomial GLM, C-start probability ∼ Amplitude + Contrast + Freezing, n = 676, for each effect p < 0.0001). For non-freezing animals C-start probability covers a range between 0.07 and 1 (mean of 0.69, Figure 4A), while in freezing animals, response probability ranges between 0 and 0.875 (mean of 0.26, Figure 4B). The effects of contrast and amplitude over C-start probability were still significant for split data (binomial GLM for non-freezing animals; amplitude: p < 0.0001; contrast: p < 0.0001; binomial GLM for freezing animals; amplitude: p < 0.0001; contrast: p = 0.003). Alarm response probability for multisensory trials ranged between 0 and 0.39, decreases with sound amplitude (p = 0.0006) but it is not affected by changes in contrast (p = 0.967, binomial GLM, Alarm response ∼ Amplitude + Contrast, n = 665, Sup. Fig. 3A).

**Figure 4.**
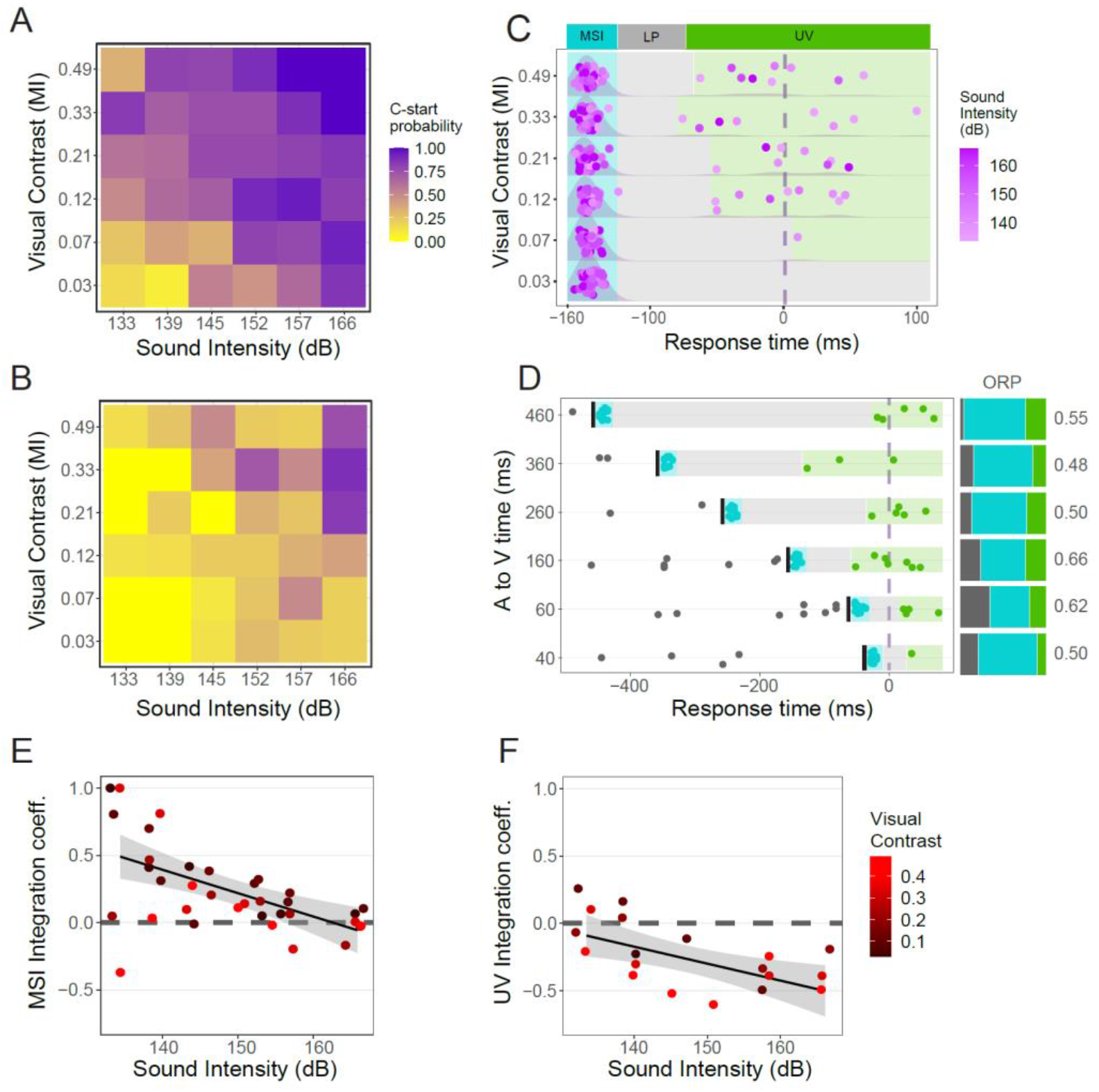
Multisensory integration enhances C-start probability and reduces response time. (A) C-start probability for non-freezing trials (n = 416) and (B) freezing trials (n = 260). (C) Response times and overlaid density distribution of C-starts during multisensory stimulation, grouped by visual contrast. Vertical scatter added for clarity. The purple shade of each dot represents the amplitude of the auditory stimulus presented 160 ms before the end of visual expansion (dashed line). The 250 ms period after the auditory stimulus shows three distinct phases: a high concentration of responses right after auditory presentation (MSI, cyan), an interval of low response probability (LP, light grey) and unisensory visual (UV) responses occurring around the end of visual expansion (green) (n = 340). (D) Multisensory response time was modulated by the delay between the auditory and visual stimuli. Sound intensity (149 dB) and visual contrast (MI 0.16) were invariant while the delay between auditory cue onset and end of loom was varied. Black bars signal the auditory stimulus. Response times follow the same pattern as in C, but the LP interval increases with the auditory-visual delay (n = 136). Stacked bars show response probability before the auditory stimulus (dark grey), in the MSI (cyan) and UV (green) periods for each delay. Numbers to the right indicate overall observed response probability (ORP). The expected response probability for this specific combination of stimulus intensities was 0.47. (E) ICs during the MSI period as a function of sound intensity colored by visual contrast (horizontal jitter applied to ease visualization). (F) ICs during the UV period as function of sound intensity and colored by visual contrast. Coefficients were calculated only for combinations which yielded responses within this period (19 out of 36). In E and F the horizontal dashed line represents no integration and the solid line and shaded area represent a linear fit to the data and 95% confidence intervals.

### Multisensory integration reduces response time

Timing is critical during a predatory evasion where a delayed escape can mean death. In this context, any additional process that speeds up threat assessment would become functionally advantageous (4). We thus asked if C-start response time was reduced in multisensory vs. unisensory conditions. Figure 4C shows response times corresponding to the same 36 multisensory combinations grouped by Michelson contrast and colored by sound amplitude. We excluded responses which occurred before auditory stimulus onset as these are unisensory visual responses (15% of the total).

In sharp contrast with unisensory experiments where density distributions of response time showed a single peak (Figure 3E-F) multisensory experiments have a more complex structure of response times with three distinct temporal windows. There is a high concentration of responses (between 82 and 100%, depending on the loom contrast) during the first 40 ms after auditory stimulus onset, which we consider true multisensory escapes and thus named this period the Multisensory Integration interval (MSI). This is followed by a low response probability window (LP) of at least 45 ms where response probability drops to 0. Finally, a small proportion of the responses (10% of the total) occur on a +/- 80 ms period centered on the end of the visual expansion (unisensory visual window, UV). Most of these late UV responses were produced when fish experienced a multisensory stimulus where the loom was combined with a low intensity sound. In contrast, multisensory stimuli that evoked responses during the MSI period had a mid to high intensity for the sound component (Supp. Fig. 3B). These results clearly show that the addition of an extremely brief sound pip is enough to markedly shift the response distribution, effectively decreasing response time compared to visual only conditions (3,30). In addition, multisensory responses latencies are much less variable than their unimodal visual counterparts, instead of spreading during more than a second, most responses are locked to the occurrence of the auditory pip (31).

To explicitly test if the onset of the LP period was dependent on the moment of auditory stimulus presentation, we performed an additional experiment where the delay between the auditory pip and the end of the visual expansion was systematically varied (−40, -60, -160, -260, -360 and -460 ms relative to the end of visual expansion) while keeping loom contrast and sound intensity constant at an intermediate value (Michelson index of 0.16 and sound amplitude of 149 dB). Figure 4D shows the response times for each of these delays. Although in some trials fish escaped before the auditory stimulus onset (i.e. before the proper multisensory event starts, grey dots, Figure 4D), most responses were triggered following the auditory stimulus, within the MSI period (cyan dots) while a small proportion of late responses were observed towards the end of the expansion (green dots). The proportion of early, MSI and UV responses for each delay is represented in the stack bars of figure 4D. Interestingly, the duration of the LP period (grey area) was not fixed but dependent on the delay of the auditory pip, with longer delays producing longer LP periods.

This particular combination of loom contrast and sound intensity produces up to a 40% enhancement in observed response probability (ORP) compared to the expected (ERP) probability (ORPs ranges between 0.48-0.66 vs. ERP=0.47). Varying the delay between the auditory pulse and the end of the expansion does not change response probability (binomial GLM, p = 0.28) nor the proportion of responses occurring during MSI (binomial GLM, p = 0.51, cyan area of stacked bars, Figure 4D). Overall, the results suggest that a relatively weak sound stimulus occurring up to half a second before the end of the visual expansion significantly reduces C-start response delay while enhancing response probability. Conversely, while addition of visual information does not further narrow the response distribution of auditory latencies it does enhance response probability.

### Multisensory integration enhances response probability

Although C-starts concentrated within the MSI, it could be that animals were either responding just to the loom or just to sound thus not necessarily implying integration. To infer actual multisensory integration, we compared response probabilities in multisensory trials with those observed during unisensory trials. For each combination of stimuli, we calculated an Integration Coefficient (IC) for the MSI period based on the ORP and the ERPs assuming no integration (see Methods). Figure 4E shows ICs plotted as a function of sound intensity during the MSI window for each of the 36 multisensory combinations tested (or as a 2D matrix, Supp. Fig. 3C and plotted as a function of visual contrast in Supp. Fig. 3D). Most ICs (28/36) during the MSI period are positive and significantly different from 0 (One sample T-test, p = 0.0002), implying that there is an overall multisensory enhancement of the C-start response, i.e. the response probability exceeds what is predicted for independent processing of unisensory cues. ICs show a negative relationship both with sound amplitude (Fig. 4E, p = 0.0001) and visual contrast (Supp. Fig. 3D, p = 0.03) although the latter effect is weaker. This negative relationship implies that audiovisual multisensory integration shows inverse effectiveness with stimulus salience. In other words, multisensory enhancement is maximized when a weak auditory cue combines with a low contrast loom (Supp. Fig. 3C) but the enhancement vanishes as unisensory cues grow in saliency (30–32).

We calculated the same coefficients for the UV period (Figure 4F and Supp. Fig. 3E). ORP during the UV period was in most cases lower than the ERP yielding negative coefficients (15/19 coefficients lower than 0) and significantly different from 0 (One sample T-test, p = 0.0002). Therefore, C-start probability during the UV period is less than what would be expected if no integration occurs, and even lower than expected for unimodal visual stimulation. We hypothesize that this reduction in the ICs during UV window is explained by the auditory stimulus anticipating responses to the MSI period. In fact, stronger sound stimuli should be more likely to anticipate responses to the MSI period (Supp. Fig. 3B), decreasing the number of responses observed in the LP and UV windows. As goldfish do not perform multiple C-starts during a trial, if an animal C-starts during the MSI, it won’t do it later during the LP and UV periods. Indeed, a Gaussian GLM tested the dependence of ICs with respect to stimulus intensity (IC ∼ Amplitude + Contrast) in the UV period. ICs show a negative relationship with sound amplitude (p = 0.008, Figure 4F) and visual contrast (p = 0.015, Supp. Fig. 3E).

Overall, these results confirm that a brief auditory cue combined with a low contrast visual loom enhances escape responsiveness compared to unisensory stimulation and reduces response time. However, this enhancement is tightly restricted to a 40 ms window following the presentation of the auditory pip. When responses reappear after the LP period, ICs show negative integration. These results prompted the question of the mechanism underlying the auditory effect on visual processing to effectively decrease response time.

### An integrate-and-fire neuron model explains multisensory enhancement of C-start behavior

In teleost fish, a variety of motor networks have been implicated in initiation and execution of evasive maneuvers (17,22,33,34). However, initiation fast-start escape responses in response to auditory pips or fast looms is in most cases triggered by a single action potential of the Mauthner cell (17,20,22,29,35). The Mauthner cell receives excitatory visual input from the tectum and auditory input from the 8th nerve in addition to shunting feedforward inhibition (36,37). Both excitatory and inhibitory inputs are integrated at the soma and if inputs are strong enough an action potential that initiates the C-start is fired. To explore the mechanism underlying the multisensory integration observed experimentally, we implemented a Leaky Integrate and Fire unit to produce a simplified model of the Mauthner cell (38,39). Here we did not attempt to produce a detailed model of the Mauthner cell circuit, but to determine the minimal parameters that could reproduce our empirical data.

Brief square pulses or ramps of increasing current fed into the model neuron represented the excitatory component of the auditory and visual inputs, respectively (Figure 5A). The intensity of six “visual” and six “auditory” current inputs were adjusted to obtain a unisensory C-start probability for each stimulus that matched the response probability that was observed empirically (Figure 5Bi and ii; compare with Figure 3A-B). As in the behavioral experiments, in the model cell C-start probability increases both with auditory intensity (binomial GLM, p < 0.0001) and visual intensity (binomial GLM, p < 0.0001). Next, we ran the simulation for each of the 36 combinations of multimodal stimuli introducing a fixed delay of 160 ms between the auditory input and the end of the visual input to obtain multisensory response probabilities (Figure 5C). Increasing auditory or visual intensity produced a significant increase of response probability (binomial GLM, C-start prob. ∼ Amplitude + Contrast, p < 0.0001 for both effects). Comparing each combination’s probability with the corresponding probability from empirical data, we find that although the model produces slightly higher C-start probabilities for the least intense combinations (C-start probability ranges between 0.35 and 0.95), the overall difference is not significant, matching the experimental results (Paired Samples T-test, p = 0.47, Figure 5C).

**Figure 5.**
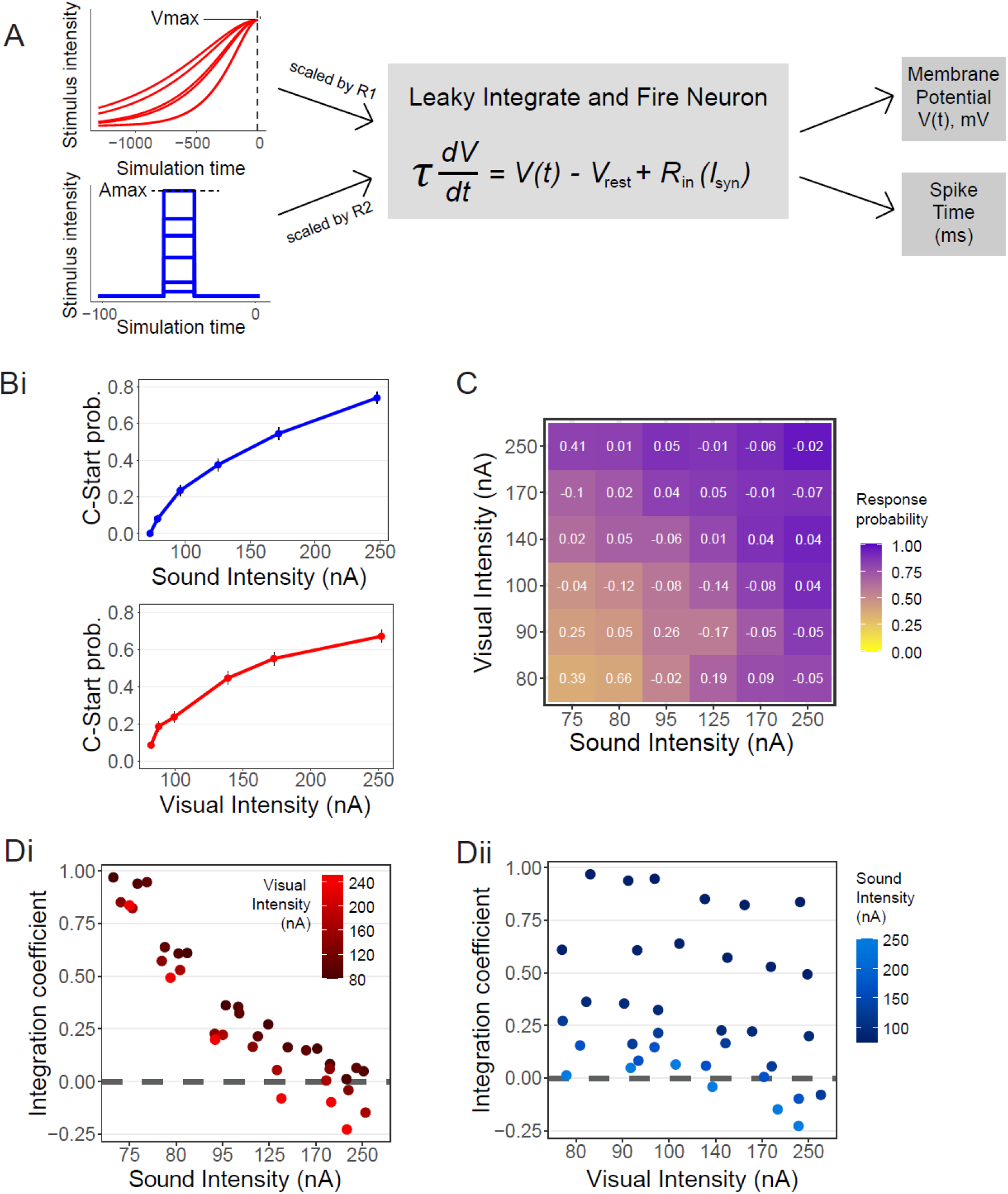
An integrate-and-fire neuron model explains multisensory enhancement of C-start behavior. (A) Model Diagram. Functions representing visual (upper left) or auditory (lower left) input converge onto the Mauthner cell model (central panel). Output variables measured are membrane potential and spike time when threshold is reached. (B) Response probability as a function of stimulus intensity for unisensory trials. Six visual (top) or auditory (bottom) intensities were fitted so that response probability matched behavioral results for non-freezing animals (n = 1200). (C) C-start probability for multisensory trials. Numbers in white represent the absolute difference with the experimental response probability (cf., Fig. 5A). (D) ICs as a function of sound intensity and colored by visual intensity (i) or as function of visual intensity and colored by auditory intensity (ii). As in behavioral experiments, the model shows enhanced multisensory integration during the MSI period and inverse effectiveness for auditory intensity, although not for visual intensity. The dashed horizontal line represents no integration.

ICs calculated within the MSI period are positive (One sample T-test, p < 0.0001) and decrease with auditory (Figure 5Di, binomial GLM, p < 0.0001) but not with visual intensity (Figure 5Dii, binomial GLM, p < 0.357), paralleling the inverse effectiveness observed for experimental data. Also, as in experimental data, ICs computed for the late window (i.e. those which occurred after the MSI) are negative (One sample T-test, p < 0.0001, not shown).The agreement between modelled and experimental data reveals that inverse effectiveness can be explained by solely considering the summation of excitatory visual and auditory signals in the Mauthner Cell.

We further asked if the model reproduced the temporal distribution of responses observed in behavioral experiments. Specifically we asked if the introduction of the “auditory” square pulse during the ramped “visual” input shifted the moment the model cell crossed threshold (i.e. the cell fired). Figure 6A shows response times for the six intensities of unisensory visual stimuli used. The density distributions peak towards the end of the stimulus (at time 0) while higher intensities produce a higher proportion of early responses, i.e. the distribution tails to the left. Both effects, as well as the range of the model response times, matched those observed in the behavioral data (Figure 3E).

**Figure 6.**
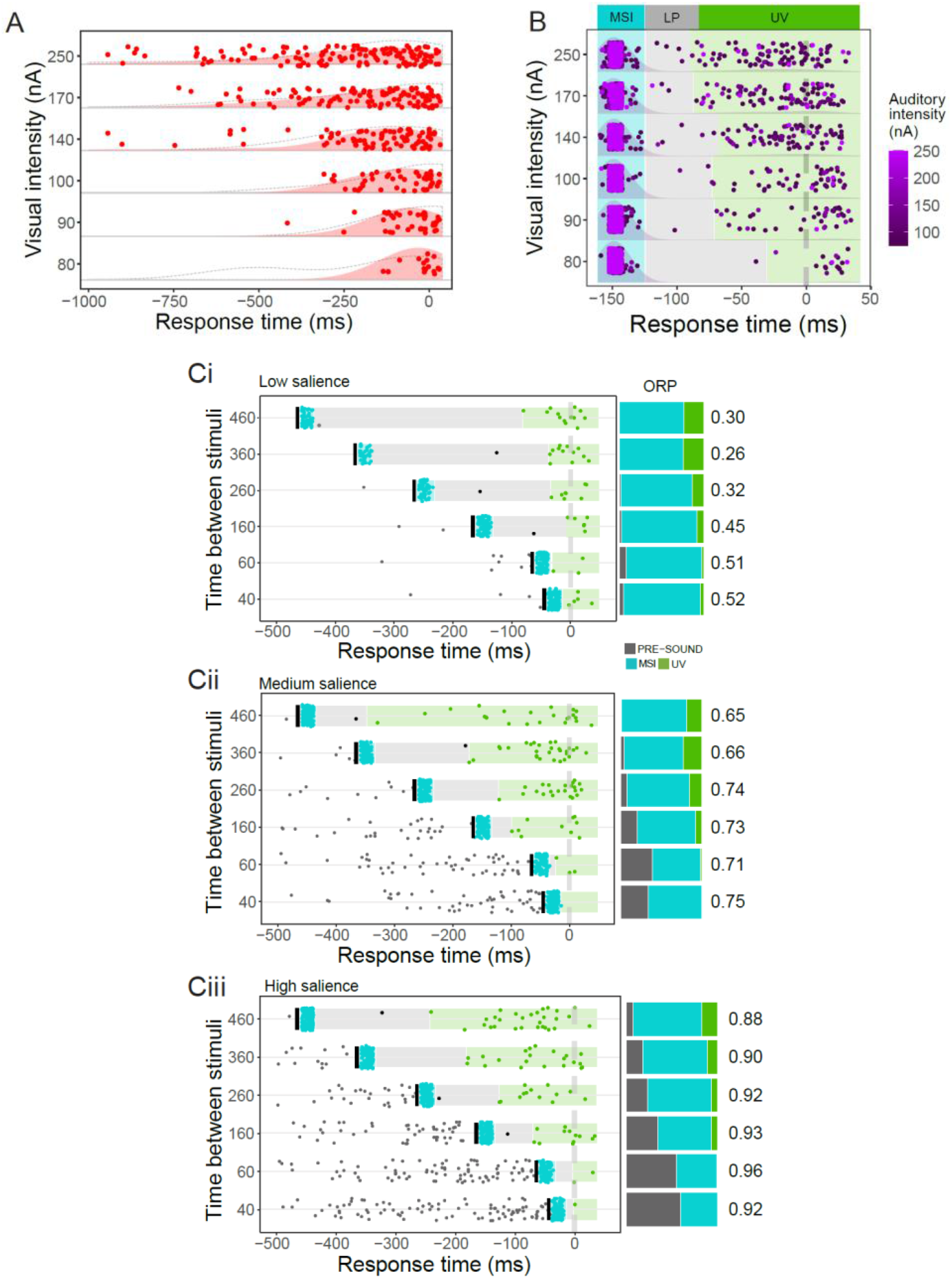
Model reproduces empirical response time distribution. (A) Response times and overlaid density distribution (light red) for visual unisensory trials, grouped by visual intensity. The grey dotted line is the experimental density distribution shown for comparison (cf. Fig. 4A). (B) Multisensory response times and overlaid density distribution, grouped by visual intensity. The color of each dot represents the auditory intensity. The vertical dotted line represents the end of visual stimulation. Responses are distributed in three distinct periods as in Figure 6 (n = 5089). The boundary between LP and UV intervals is set by 5 and 95% of the responses. (C) Response times for multisensory trials with variable delay between the auditory stimulus and the end of visual stimulus for low (i) medium (ii) or high (iii) salience. Black vertical segments represent the time of auditory stimulus presentation (n = 845). Stacked bars to the right show response probability before the auditory stimulus (dark grey), MSI (cyan) and UV (green) for each delay. Numbers to the right indicate overall observed response probability for the simulation (sORP). Simulation was ran 100 times for each condition.

As in empirical data, simulated multisensory responses after auditory stimulus onset were divided into three distinct periods (Figure 6B). A high concentration of responses (between 82% and 97% depending on the contrast) occurred during the MSI period. This was followed by a sharp decrease in response probability during the LP period, which got progressively longer as visual intensity decreased. Finally, a relatively low number of event occupied the UV period (compare with figure 4C). The model (run ten times more than behavioral trials) shows robustly that response probability in the LP and UV windows is not fixed but it increases for higher visual intensities, a trend that was present but subtle in the behavioral experiments. In also shows that most UV responses correspond to stimuli in which the sound component was weak.

We next explored the effect of systematically changing the delay between the auditory and visual inputs at three different saliency levels. Figure 6Ci-iii shows response times for combinations of auditory and visual stimuli of low, medium and high salience, where the elapsed time between auditory stimulus onset and the end of visual expansion was varied as in figure 4D. The simulation reproduced the empirical temporal structure of response times (compare Figure 6Cii with Figure 4D). It also made evident that as visual saliency increases, the proportion of early (visual unimodal) responses grows (grey stacked bars, Fig. 6Ci-iii). While behaviorally we tested a single combination of intermediate sound intensity and visual contrast with different delays, modelling shows that comparable results are obtained when salience is lower (Figure 6Ci) or higher (Figure 6Ciii). Taken together, computational results strongly suggest that that the multisensory enhancement of the C-start response and its inverse effectiveness features can be explained by only considering the interaction of excitatory visual and auditory signals in the Mauthner Cell. Moreover, the temporal structure of the C-start response distribution also can be explained by a model that simply accumulates excitation until reaching threshold.

## DISCUSSION

The main questions posed in this paper are how goldfish integrate sensory information during risk assessment and whether this varies with the salience of the multisensory stimulus. We found that the addition of a brief sound pulse is capable of enhancing detection while speeding up the response to a visual threat. Multisensory enhancement disappears as unimodal saliency increases to make single stimuli strong enough to bring the Mauthner cell to threshold. Providing mechanistic grounds for these observations, we found that behavioral results are reproduced by an Integrate and Fire model neuron. Noticeably, it was enough to combine excitatory input currents with dynamics matching the temporal structure of the empirical auditory and visual stimuli to reproduce the multisensory response enhancement and the reduction in response time.

### Inverse effectiveness of multisensory stimuli is strongly dependent on auditory intensity

Hallmark of multisensory integration, inverse effectiveness is traditionally explained in terms of the ambiguity of the situation (1,2). When any component of a multisensory percept is very salient, additional information contributed by secondary components will make only a modest contribution to the response. However, when unisensory stimulus strength is lowered, the value of combining information for independent sources grows. Although we matched the range of saliences of our visual and auditory stimuli (using response probability as proxy for salience), increasing sound intensity produced a steeper decrease in multisensory enhancement than increasing contrast (i.e. a stronger inverse effectiveness, compare fit slope in Fig. 4E vs. Supp. Fig. 3D). From a functional perspective, one could argue that a blunt noise is a much less ambiguous signal of immediate danger than a gradual size increase of a dark spot. More mechanistically, considering the neural substrate of the C-start, a simplified model of the Mauthner cell circuit reproduces the asymmetry on the inverse effectiveness of visual and auditory salience (Fig. 5Di vs. ii). This model consists of a single compartment that integrates excitatory currents (Fig. 5A), and thus all presynaptic circuit effects as well as dendritic filtering that may vary between auditory and visual processing are omitted (13). The only difference remaining between the “visual” and “auditory” components is their temporal structure. It is therefore not necessary to postulate complex differences between auditory and visual processing (even though they may exist) to explain their impact in response probability, multisensory integration and inverse effectiveness: in this case, it suffices to account for distinct temporal dynamics. This underscores that the temporal structure of a sensory stimuli adds critical information to its meaning.

### Reduction in response time to multisensory stimuli is determined by sound onset

The auditory stimulus not only had a relatively stronger inverse effectiveness but also definitive influence on response time. The temporal distribution of unisensory visual responses peaks towards the end of the loom but has a tail of earlier responses which gets increasingly skewed as contrast increases (Fig. 3E and 6A). When a sound pip is added even as early as 460ms before the end of the expansion most responses concentrate in a narrow MSI time window following the pip. This effect is observed irrespective of loom contrast, sound intensity (Fig. 4C and 6B) or the interval between the sound and the end of the expansion (Fig. 4D and 6C). However, the MSI period is not just a response to an acoustic stimulus. The width of the response time distribution for multisensory trials doubles the duration of the auditory unisensory responses (40 vs. 20 ms, Fig. 4C vs. 3F). Functionally, adding a brief auditory pip to a low contrast looming expansion increases response probability and reduces reaction time compared to a visual only condition. In a real-life predatory encounter the importance of binding this two pieces of ambiguous information becomes evident (4).

Following the MSI period we consistently observed a time window where no responses were recorded (LP) and where multisensory integration becomes negative. Auditory-evoked feedforward inhibition of the Mauthner cell could be implicated in the lack of responses during the LP period, but inhibition decays after a few tens of milliseconds (15). Although we cannot rule out that inhibition is delaying the UV responses, our computational model was able to reproduce the pattern of an MSI period followed by a LP window without having specified such a mechanism. Conceptually, cells that are sufficiently depolarized by the loom to fire before the end of expansion will cross threshold the moment a sudden auditory stimulus arrives, providing the auditory amplitude is strong enough (red trace, Figure 7). Less depolarized cells that fail to fire when the auditory stimulus is paired with the loom will fire, if at all, near the end of visual expansion, when visual excitatory input is strongest (pink trace). Finally, cells which are more excited or that receive a stronger visual input will reach threshold before the sound stimulus occurs (dark red trace). Although additional neural circuitry surely play a role in multisensory integration and escape execution, our findings suggests that excitatory inputs integrated at the Mauthner cell soma are key elements in multisensory decision making during fast C-start escapes.

**Figure 7.**
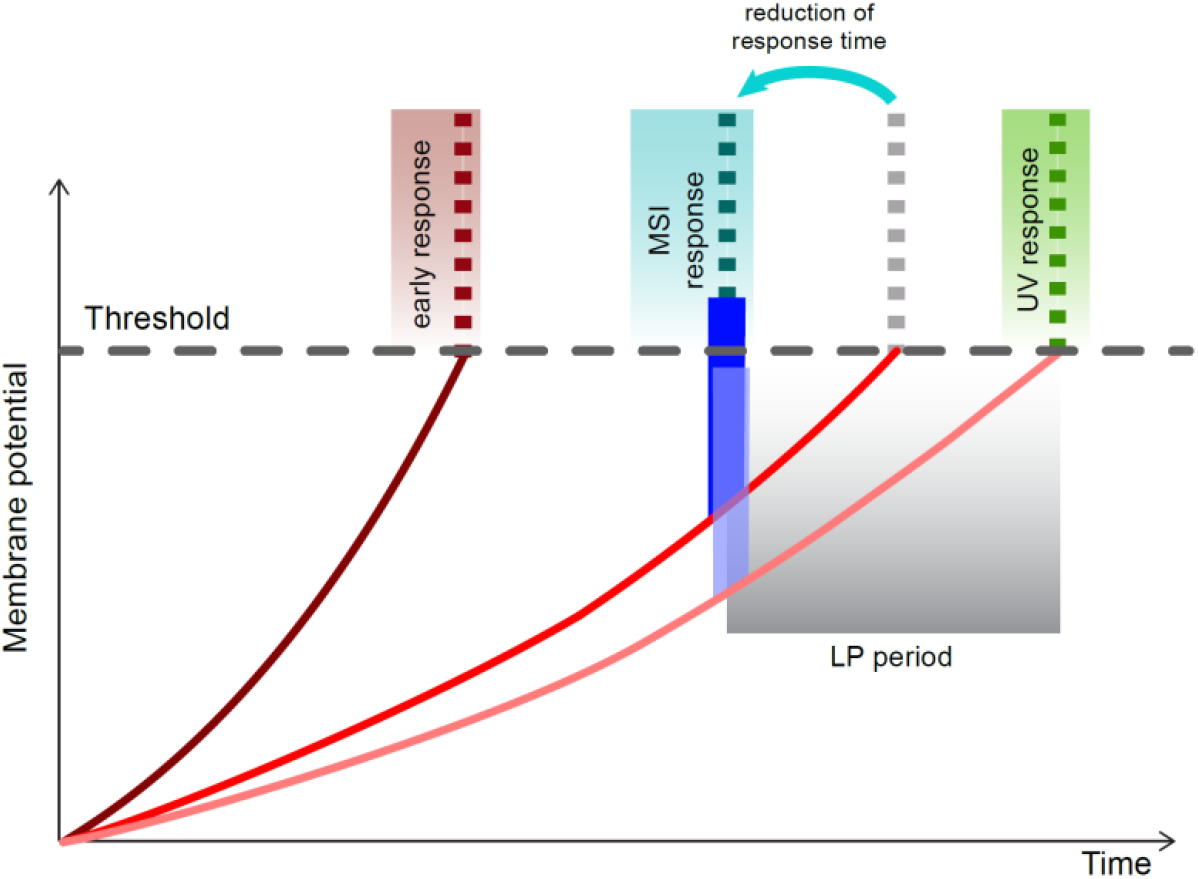
Conceptual model of multisensory integration in the Mauthner cell. Mauthner cells showed a ramped depolarization as a consequence of the increasing visual expansion (red curves). Intrinsic differences in cell excitability or salience of the stimulus could produce different rates of depolarization. As a consequence of the sound pulse (blue rectangles) there is a sharp and brief increase in membrane potential a few milliseconds after sound onset. If membrane potential surpasses the firing threshold (black dotted line), a C-start response is evoked (dotted lines). When the system depolarizes at a higher rate (dark red), early (unisensory visual) responses occur. Lower excitability (lightest red) results in late (or no) responses, since auditory stimulation is not enough to reach threshold (light blue pulses). Intermediate excitability results in audiovisual C-starts locked to the time of sound presentation if the auditory response mounted on the visually evoked depolarization reaches threshold (red). As a consequence, multisensory integration decreases response time and produces a low response (LP) period where unimodal visual stimuli with intermediate excitability would produce responses in the absence of a sound stimulus.

## SUPPLEMENTARY FIGURES

**Supplementary Figure 1.**
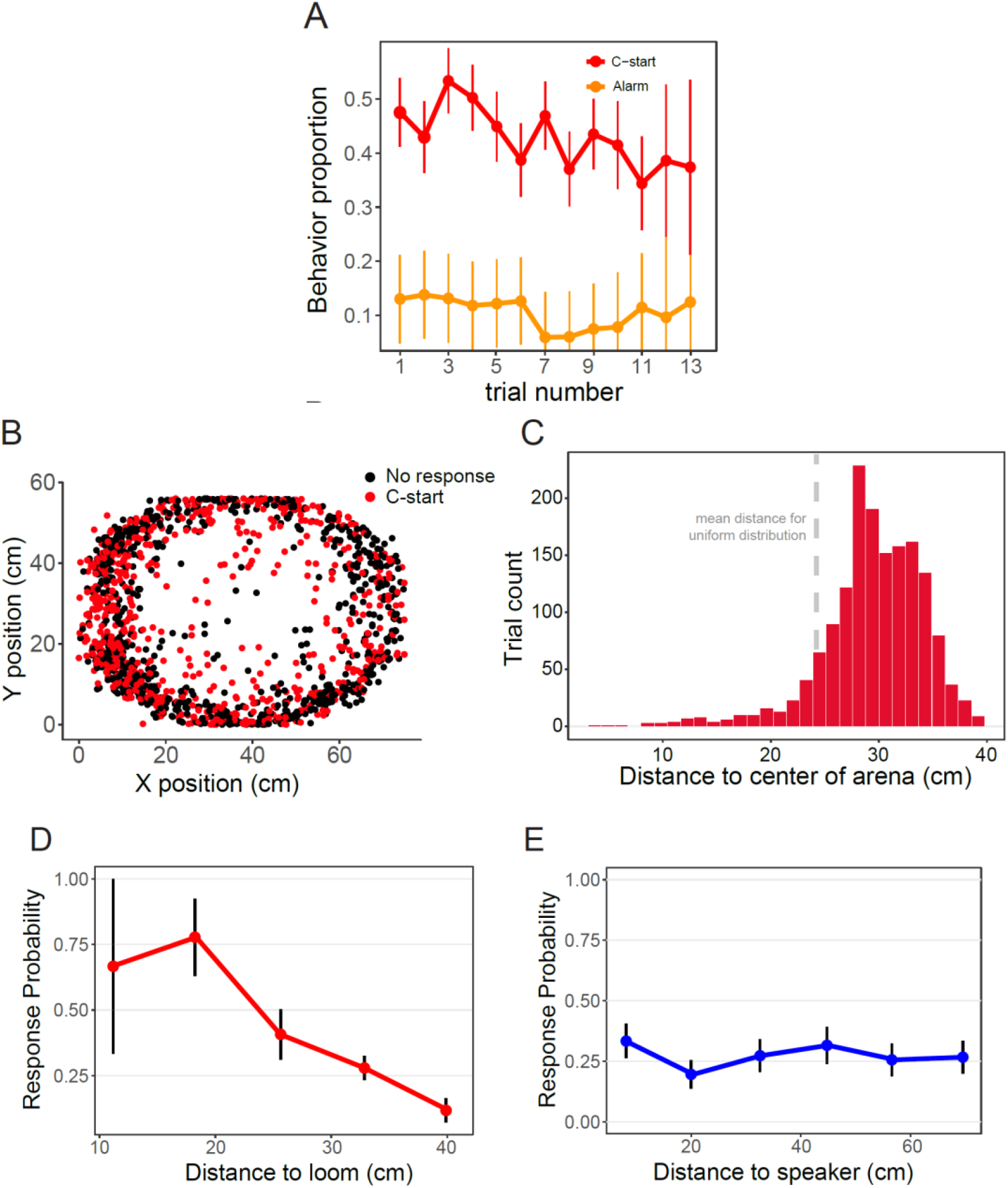
Influence of order of stimulus presentation and fish position in experimental arena. (A) Response probability decreases as the number of stimulations increases, especially for C-start responses but also slightly for alarm responses. Data from all behavioral experiments combined (n=1419, N=141). Error bars represent standard error for the proportion. (B) Position in experimental arena is not homogenous, a strong thigmotactic behavior is observed. Each dot represents the position of a fish in a single trial, at the time of C-start response (red dots), or 1500 ms before the end of visual expansion for non-responding animals (black dots). Data from all behavioral experiments combined. (C) Distance from fish head to the center of the experimental arena is not uniform. The distribution of distances is significantly shifted towards larger distances, confirming the thigmotactic behavior. Dotted gray line represents the mean of a theoretical uniform distribution. (D) Response probability decreases as distance to loom increases. Data from all unisensory visual stimuli combined (n=183, N=80). (E) Response probability is not dependent on distance to speaker. Data from all unisensory auditory stimuli combined. (n=261, N=106). In A, D and E black bars show binomial standard error.

**Supplementary Figure 2.**
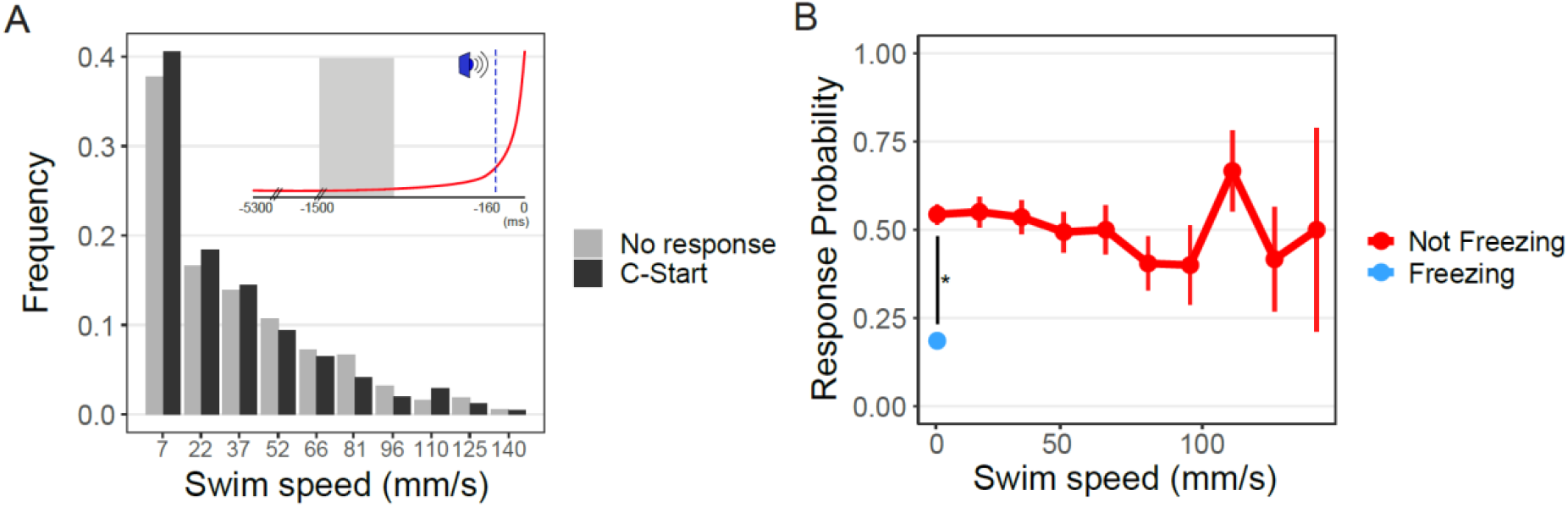
(A) Frequency distribution of swimming speeds prior to stimulation is similar in trials which ended in C-start or not (n C-start = 414; n No-response = 374). Inset shows the time interval (grey shading) where swim velocity was measured. (B) Response probability vs. swimming speed for not freezing (red) compared to freezing animals (light blue). Although C-start response probability does not depend on swimming speed it is significantly higher for still (zero velocity) than truly freezing animals.

**Supplementary Figure 3.**
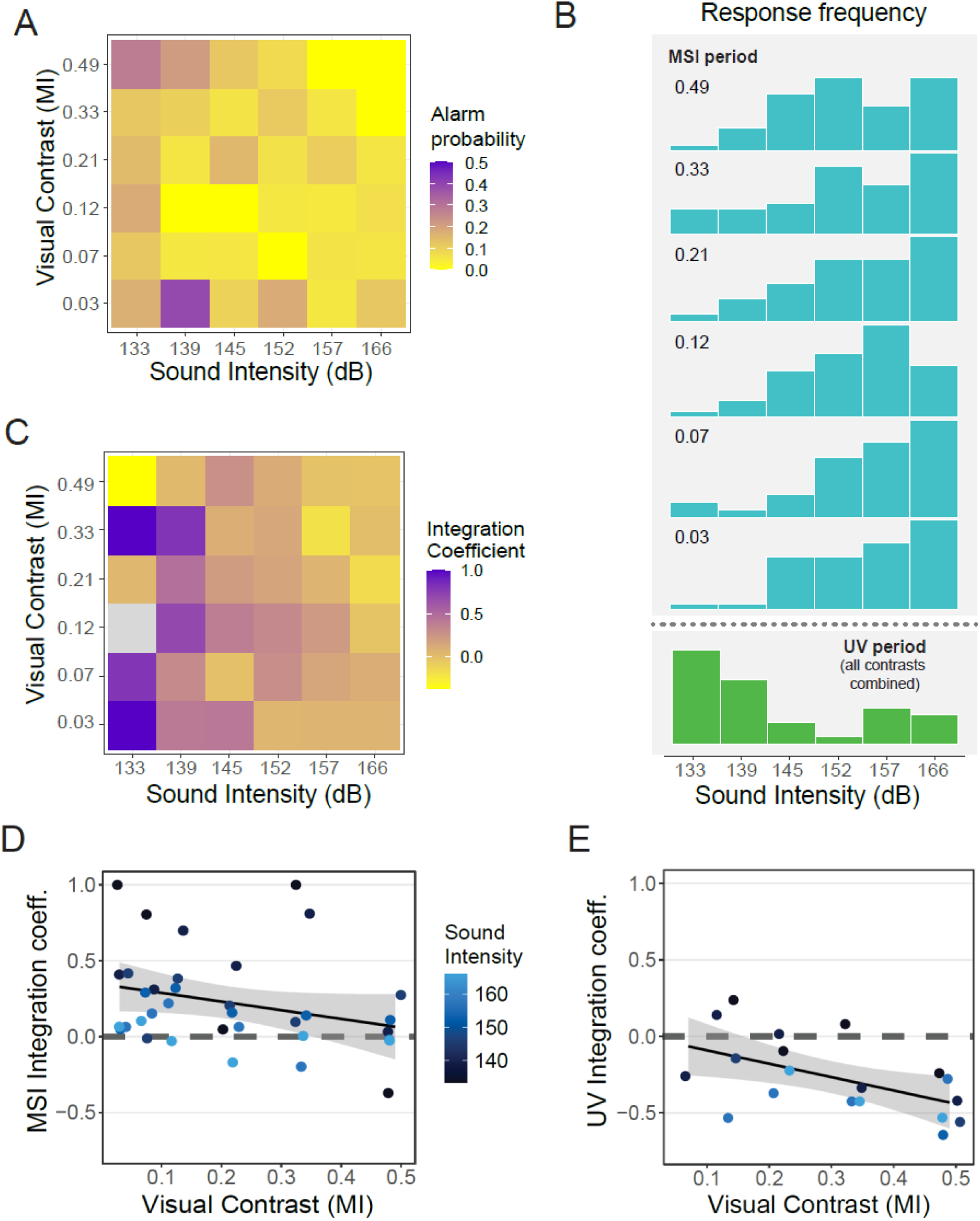
(A) Alarm response probability for each multisensory combination (n = 664). Note the intensity scale runs from 0 to 0.5%. Since 93% of all alarm responses were performed by non-freezing animals, data was not split. (B) Histograms of the intensity of the sound component of responses on MSI period sorted by visual contrast (numbers in upper left corner, cyan) or during the UV period (green). In the latter, all contrasts where combined due to the low number of observations. (C) ICs during the MSI period for each combination of visual contrast and sound intensity. There were no responses for the combination shown in grey. (D) ICs during the MSI period as a function of visual contrast colored by auditory intensity (horizontal jitter is applied to ease visualization). (E) ICs during the UV period as function of visual contrast colored by auditory intensity.

